# Low-salt diet induces claudin-3 expression and drives adaptive changes in collecting duct of claudin-3-deficient mice

**DOI:** 10.64898/2026.01.26.701749

**Authors:** Ali Sassi, Alexandra Chassot, Sara Jellali, Nicolas Liaudet, Ana Polat, Felix Baier, Deborah Stroka, Mikio Furuse, Eric Feraille

## Abstract

**Background:** Renal sodium reabsorption occurs *via* both transcellular and paracellular pathways. Tight junction proteins play a key role in mediating paracellular transport. The collecting duct (CD) is critical for the fine-tuning of Na^+^ balance and is sensitive to changes in dietary salt intake. A low-sodium diet, which increases endogenous aldosterone secretion, stimulates transcellular sodium transport *via* epithelial Na^+^ channels (ENaC) and Na,K-ATPase. We hypothesized that a low-sodium diet also modulates paracellular Na^+^ permeability by regulating the expression or function of claudin-3, a major tight junction protein in the CD, in order to limit the back-leak of reabsorbed sodium and preserve sodium balance.

**Methods:** We used *in vivo* mouse models and cultured mouse CD principal cells (mCCD_cl1_) to assess aldosterone’s effects on tight junction proteins. In mCCD_cl1_ cells, aldosterone-induced changes in claudin-3 expression and localization were evaluated via Western blotting and immunofluorescence, and Ussing chamber assays were used to assess paracellular Na^+^ and Cl^−^ permeability after modulating claudin-3 expression. Wild-type and claudin-3 knockout mice were fed low (0.01%) or normal (0.18%) sodium diets for seven days. In subsets of low sodium diet mice, spironolactone (a mineralocorticoid receptor antagonist) was administered.

**Results:** In mice, a low-sodium diet upregulates renal claudin-3 expression. Concordantly, *in vitro* studies using mCCD_cl1_ cells showed that aldosterone treatment increased claudin-3 protein levels and promoted its localization to the lateral membrane. Functional analyses demonstrated that claudin-3 overexpression reduced paracellular permeability to both Na^+^ and Cl^−^, while claudin-3 silencing increased it. Claudin-3 knockout mice subjected to a low-sodium diet exhibited compensatory upregulation of the α- and γ-subunits of ENaC, alongside increased expression of claudin-4, claudin-8, and claudin-10. This highlights an adaptive response that maintains sodium homeostasis in the absence of claudin-3. Importantly, this compensatory mechanism persists even under spironolactone treatment, suggesting that the adaptation of claudin-3-deficient mice occurs independently of mineralocorticoid receptor activation.

**Conclusions:** Our findings demonstrate that aldosterone enhances claudin-3 expression, reinforcing the paracellular barrier to Na^+^ and complementing its classical role in transcellular Na^+^ transport. Under low-sodium conditions, claudin-3-deficient mice adapt through complementary mechanisms aimed at increasing sodium reabsorption *via* ENaC activation and upregulation of claudin-4 and claudin-8, both barrier-forming claudins that restrict paracellular sodium leakage in the CD. This is associated with increased claudin-10 abundance in the thick ascending limb of Henle, a pore-forming claudin that facilitates paracellular sodium permeability. This study advances our understanding of the complex control of renal sodium handling, revealing adaptive mechanisms in response to low-salt diet and claudin-3 deficiency.

## Introduction

The regulation of sodium (Na^+^) transport by the kidney is essential for maintaining electrolyte balance, fluid homeostasis, and blood pressure. The collecting duct (CD), the final site of regulated Na^+^ reabsorption along the kidney tubule, plays a crucial role in this process. Sodium transport in the CD occurs primarily through the epithelial sodium channel (ENaC) located at the apical membrane and the Na,K-ATPase located at the basolateral membrane, the latter generating the electrochemical gradient necessary for transcellular Na^+^ reabsorption (1, 2). In addition to transcellular transport, paracellular transport is highly dependent on tight junctions (TJ), which may generate either a specific Na^+^ permeability or a barrier that prevents passive Na^+^ back-leak into the tubular lumen and thereby maintain a net Na^+^ reabsorption process (3). TJ are composed of various proteins, including occludin, junctional adhesion molecules (JAMs), and claudins. Claudins are the primary determinants of TJ selectivity, regulating the permeability of ions and water across epithelial barriers (4, 5). The expression profile of claudins in the kidney is highly segment-specific, with distinct claudins contributing to either barrier function or selective ion permeability (6, 7).

Aldosterone, the main mineralocorticoid hormone, is a key regulator of Na^+^ balance and renal sodium reabsorption. In CD principal cells, binding of aldosterone to the mineralocorticoid receptor (MR) triggers genomic and non-genomic responses that enhance Na^+^ transport (8–10). Aldosterone increases the expression and activity of both ENaC and Na,K-ATPase, thereby enhancing transcellular Na^+^ reabsorption (11–14). However, aldosterone-induced Na^+^ reabsorption generating a lumen-negative transepithelial potential, together with the Na^+^ gradient between the interstitial space and the lumen, may drive a Na^+^ back-leak through the paracellular pathway. Increased paracellular Na^+^ back-leak potentially compromises the efficiency of Na^+^ reabsorption. Aldosterone has been reported to strengthen the paracellular barrier by modulating TJ composition. For example, it increases claudin-8 expression in the kidney and colon, reinforcing the Na^+^ diffusion barrier. However, the extent to which aldosterone regulates the amounts of other claudin species along the kidney tubule TJ remains to be fully established. Claudin-3 is widely expressed in epithelial tissues, including the respiratory, urinary and gastrointestinal tracts, as well as the liver, salivary and mammary glands. Its function remains controversial: some studies suggest that claudin-3 enhances barrier function by reducing paracellular permeability, while others suggest that it may facilitate selective permeability pathways depending on the tissue and physiological context (15–21). The function of claudin-3 has been well studied in several tissues, while its specific role in the kidney remains poorly understood and requires further investigation.

In this study, we investigated the effect of aldosterone on claudin-3 expression in CD principal cells. Using *in vivo* and *in vitro* models, we demonstrated that aldosterone upregulates claudin-3 protein abundance, suggesting a potential role for claudin-3 in the hormonal regulation of paracellular Na^+^ transport. Furthermore, we examined the functional consequences of claudin-3 loss by studying claudin-3-deficient mice under a low-salt diet, which stimulates endogenous aldosterone secretion. These mice exhibited compensatory adaptations, including increased expression of α- and γ-ENaC subunits and upregulation of claudin-4, claudin-8 and claudin-10, indicating that the absence of claudin-3 triggers specific molecular adjustments to preserve Na^+^ balance under low-salt diet conditions. These findings provide new insights into the mechanisms regulating renal Na^+^ reabsorption and highlight the importance of TJ regulation in maintaining electrolyte balance.

## Methods

### Animals

For the experiments investigating the effects of low-NaCl diet on claudin-3 in wild-type mouse kidneys, male C57BL/6 mice (Charles River, Saint-Germain-de-l’Arbresle, France) were used. The generation of the C57BL/6 claudin-3-deficient mouse strain with a global claudin-3 knockout was previously described by Castro Dias *et al.,* (22). Homozygous claudin-3 knockout and Claudin3 wild-type mice were obtained by interbreeding heterozygous parents, and both the experimental (knockout) and control (wild-type) groups were bred and housed in the same facility. The mice had ad libitum access to food and water and were maintained on a 12-hour light/dark cycle. All procedures were performed during the light phase in male mice aged 8 to 12 weeks.

To assess the effect of sodium diets, mice were fed for 7 days with either a low-sodium (Na^+^) diet [0.01% (wt/wt); LSD] or a normal-sodium (Na^+^) diet [0.18% (wt/wt); NSD] (Provimi-Kliba, Kaiseraugst, Switzerland). One group of mice on the low-Na^+^ diet received 0.35 mg/100 g body weight/day of spironolactone mixed with food for 7 days.

For physiological parameter measurement, mice were acclimatized to Tecniplast metabolic cages for 24 hours. Following acclimation, food intake, water consumption, and urine output were measured over a subsequent 24-hour period.

Venous sinus blood was collected from anesthetized animals using heparinized capillary tubes to prevent clotting. Immediately after collection, the blood samples were analyzed with the epoc^®^ Blood Analysis System (Siemens Healthineers) according to the manufacturer’s protocol. This system enabled rapid and reliable measurements of key blood biological parameters.

All animal experiments were approved by the Institutional ethical committee of animal care of the university of Geneva and the cantonal authorities, in accordance with the office of laboratory animal welfare’s guidelines for good animal practice. The study also adhered to the standards set by the National Centre for the Replacement, Refinement, and Reduction of Animals in Research (NC3Rs).

### Cell culture, constructs, viral particle production, and cell transduction

As previously described, mCCD_cl1_ cells were grown on permeable filters (Transwell®, Corning Costar, Cambridge, MA) to confluence in a 1:1 mixture of Dulbecco’s modified Eagle’s medium and F12 medium, as previously described (23). Aldosterone (10^-6^ M) was added to the apical and basal compartments 24 h before cell lysis. For gene overexpression, cells were transduced with lentiviruses carrying either an empty vector or wild-type green fluorescent protein (GFP) as controls, or a construct encoding wild-type mouse claudin-3. Both claudin-3 and GFP were previously subcloned into a modified puromycin-resistant pSF-lenti vector (Sigma). For gene silencing, mouse claudin-3, or scramble short hairpin RNA (shRNA) (Table 1) was inserted into the plasmid pLKO.1 (cat. no 8453; Addgene). pLKO.1 or pSF-lenti was transiently transfected in packaging HEK293T cells using the Polyplus-transfection^®^ jetPRIME^®^ Kit according to the manufacturer’s instructions. Lentiviral particles were collected after 72 hours and mCCDcl1 cells were transduced. Stable polyclonal cell lines were selected using puromycin (2 *μ*g/ml) which was applied 72 hours after transduction.

**Table 1:**
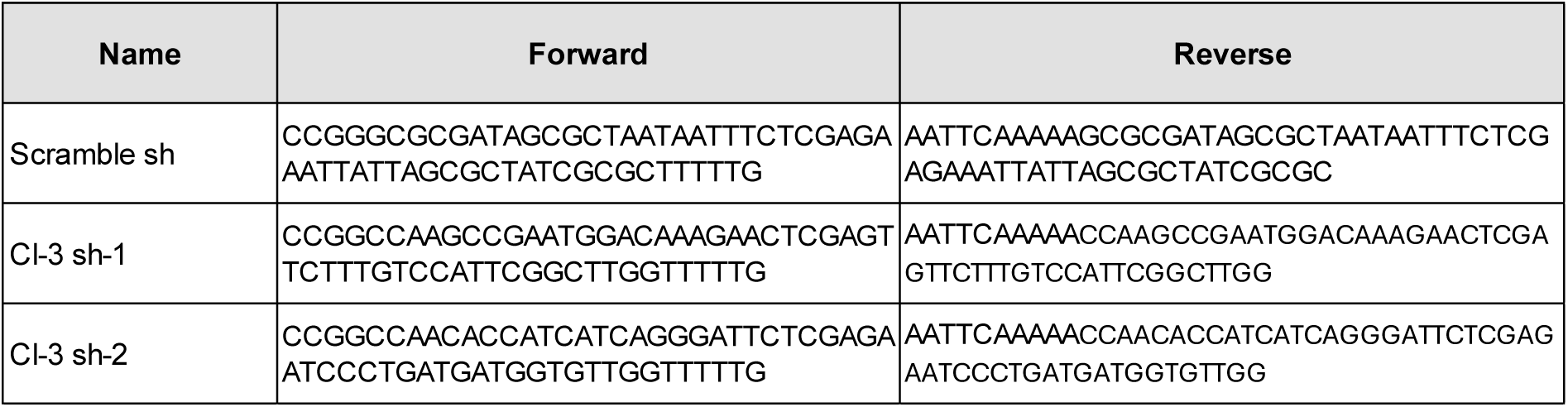
Sequences of scramble shRNA and shRNAs targeting claudin-3

### Electrophysiological assessment of paracellular permeability in mCCD_cl1_ Cell monolayers

To determine paracellular permeability, measurements were performed using cultured mCCD_cl1_ cells plated onto Snapwell polyester filters (cat. no. 3801, Corning Costar, Cambridge, MA) at a density of 250,000 cells/cm² and maintained in culture medium for seven days. Following incubation, the Snapwell inserts were detached and positioned in Ussing chambers (model P2300; Physiologic Instruments, San Diego, CA, USA). These chambers were connected to a VCC MC6 multichannel voltage/current clamp (Physiologic Instruments) using silver/silver chloride (Ag/AgCl) electrodes and 3 M KCl agar bridges.

The apical and basolateral compartments of the Ussing chambers were separately filled with buffer A (120 mM NaCl, 10 mM NaHCO_3_, 5 mM KCl, 1.2 mM CaCl_2_, 1 mM MgCl_2_, and 10 mM Hepes, pH 7.4), with each side containing a total volume of 5 ml. Transepithelial potential differences between the two compartments were recorded using a Quick Data Acquisition DI100 USB board (Physiologic Instruments), while the transepithelial current was maintained at 0 mA throughout the experiment. Prior to measurement, the cells were equilibrated in buffer A for one hour.

To determine the dilution potential for NaCl, half of the buffer in the basal chamber was replaced with buffer B (240 mM mannitol, 10 mM NaHCO_3_, 5 mM KCl, 1.2 mM CaCl_2_, 1 mM MgCl_2_, and 10 mM Hepes, pH 7.4), ensuring osmolarity balance with mannitol and adjusting pH to 7.4 with HCl. To confirm that ion flux occurred *via* the paracellular route, 100 µM amiloride (Sigma) and 100 µM 4,4′-diisothiocyanatostilbene-2,2′-disulfonic acid (DIDS; Sigma-Aldrich) were added to the apical chamber 30 minutes before measurement. Throughout the experiment, buffers were maintained at 37°C and continuously bubbled with a gas mixture of 95% oxygen and 5% carbon dioxide.

Peak dilution potential values were recorded and used to determine permeability ratios. Absolute ion permeabilities were calculated as previously described (24). Specifically, absolute permeabilities for Na^+^ and Cl^−^ were computed using the Kimizuka-Koketsu equation, incorporating both the calculated P_Na_/P_Cl_ and the transepithelial resistance measured during the experiment.

### Western blotting

Equal amounts of protein from cultured cells or kidney cortex were separated on SurePAGE™ precast Bis-Tris 4–20% gradient gel (cat. no. M00657, GenScript, Switzerland), and transferred to polyvinylidene difluoride membranes (Immobilion-P, Millipore, Bedford, MA), as previously described (23). After incubation with primary antibodies (Table 3), membranes were incubated with anti-rabbit or anti-mouse IgG antibody coupled to horseradish peroxidase (Transduction laboratories, Lexington, KY), the antigen-antibody complexes were detected by enhanced chemiluminescence (Advansta, Menlo Park, CA). Protein abundance was quantified with the image J software. Results are expressed as the ratio of the densitometry of the band of interest to the loading control.

### Immunofluorescence

Cultured mCCD_cl1_ grown on polycarbonate filters were fixed with ice-cold methanol for 5 min at -20°C and then washed with PBS during 30 min. Blocking of nonspecific binding sites was done with 10% Normal Goat Serum (NGS) diluted in Phosphate Buffered Saline (PBS). Cells were then incubated overnight at 4°C with antibodies against claudin-3 diluted 1:100 in 2% BSA followed by a 1 h incubation with Alexa Fluor 488-conjugated goat anti-rabbit (cat. no. A-11008; Invitrogen) diluted 1:1000 and were finally mounted on microscope slides using Vectashield mounting medium (Maravai Life Science, San Diego, CA, USA) with DAPI for nuclear counterstaining. Fluorescence images were acquired using a LSM 700 confocal laser-scanning microscope (Carl Zeiss, Oberkochen, Germany) using 488-nm ray lasers. The distance between the Z-slices was 0.25 μm. From 5 to10 Z-stack images were processed per sample using the ZEISS ZEN Imaging Software. ZEN 2.3 (Carl Zeiss, Oberkochen, Germany).

After dehydration and paraffin-embedding, kidneys sections were cut at a thickness of 5 μm. Antigen retrieval was done with Tris-EDTA buffer 10 mM pH 9. Permeabilization with Triton X100 0.2% in PBS followed by blocking of non-specific binding sites with 10% NGS diluted in Tris-buffered saline with 0.1% Tween 20 (TBST) were applied for 1h. The tissue sections were then incubated overnight at 4°C with primary antibodies (Table 3), in NGS-TBST followed by a 1h incubation with a secondary Alexa Fluor 488-conjugated goat anti-rabbit (cat. no. A-11017; Invitrogen) or Cyanin3-conjugated goat anti-mouse (cat. no. M30010; Invitrogen) diluted between 1:200 to 1:800 in NGS-TBST at 37 °C. Samples were mounted on microscope slides using Vectashield mounting medium (Maravai Life Science, San Diego, CA, USA). Fluorescence images were acquired using a Zeiss Axio Imager M2 (Carl Zeiss, Oberkochen, Germany). Negative controls were performed in the absence of primary antibody (not shown).

### RNA Extraction and Quantitative PCR Analysis

Total RNA was isolated from cultured cells using the EZNA Total RNA Kit I (cat. no. R6834, Omega BIO-TEK), following the protocol provided by the manufacturer. The RNA concentration and purity were assessed using a Nanodrop spectrophotometer. To generate complementary DNA (cDNA), one microgram of RNA was converted to cDNA using the qScript cDNA SuperMix (Quanta Biosciences), adhering to the manufacturer’s guidelines. The specific primers used are detailed in Table 2.

**Table 2:**
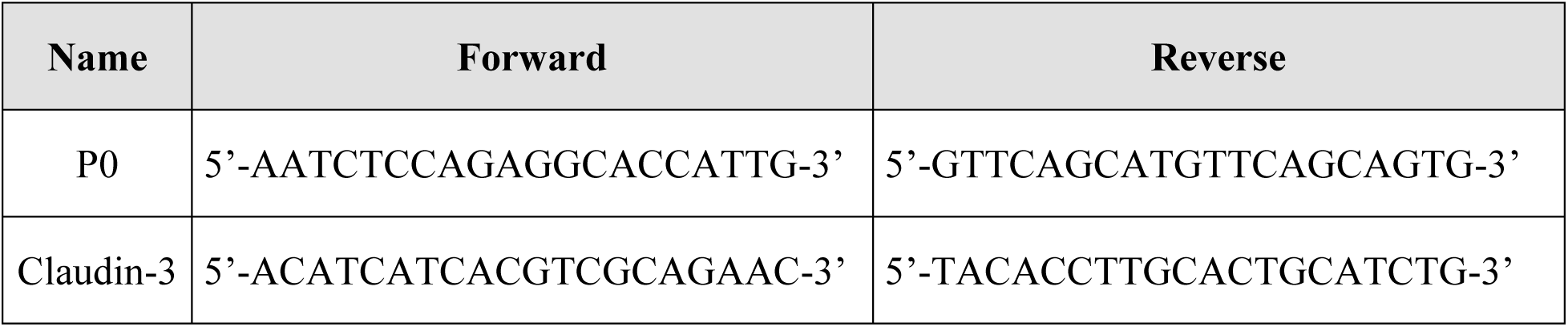
Sequences of primers used for Real-Time PCR.

**Table 3:**
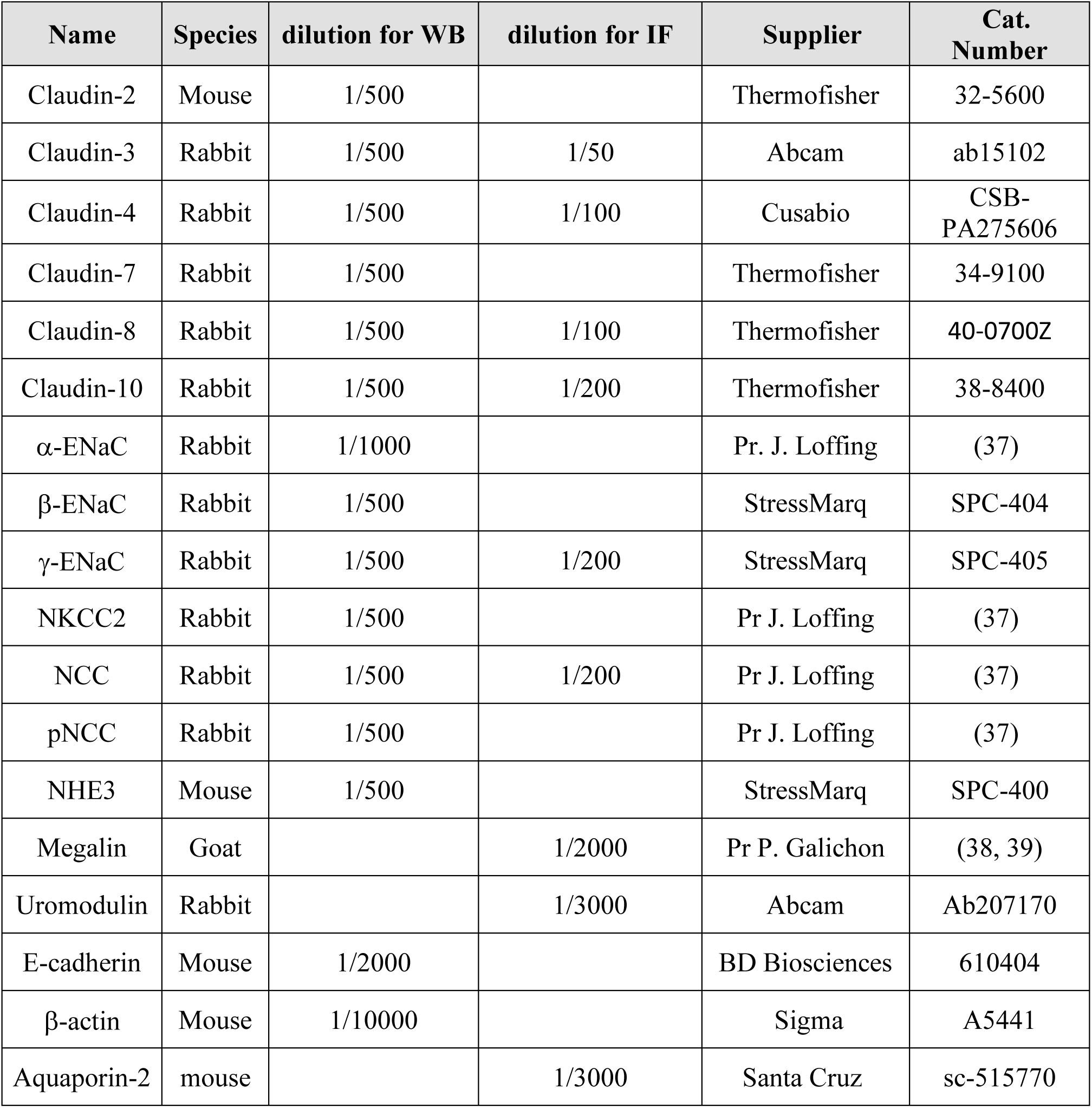
Antibodies used for Western blots (WB) and immunofluorescence (IF)

| Name | Species | dilution for WB | dilution for IF | Supplier | Cat. Number |
| --- | --- | --- | --- | --- | --- |
| Claudin-2 | Mouse | 1/500 |  | Thermofisher | 32-5600 |
| Claudin-3 | Rabbit | 1/500 | 1/50 | Abcam | ab15102 |
| Claudin-4 | Rabbit | 1/500 | 1/100 | Cusabio | CSB-PA275606 |
| Claudin-7 | Rabbit | 1/500 |  | Thermofisher | 34-9100 |
| Claudin-8 | Rabbit | 1/500 | 1/100 | Thermofisher | 40-0700Z |
| Claudin-10 | Rabbit | 1/500 | 1/200 | Thermofisher | 38-8400 |
| α-ENaC | Rabbit | 1/1000 | | Pr. J. Loffing | (37) |
| β-ENaC | Rabbit | 1/500 | | StressMarq | SPC-404 |
| γ-ENaC | Rabbit | 1/500 | 1/200 | StressMarq | SPC-405 |
| NKCC2 | Rabbit | 1/500 |  | Pr J. Loffing | (37) |
| NCC | Rabbit | 1/500 | 1/200 | Pr J. Loffing | (37) |
| pNCC | Rabbit | 1/500 |  | Pr J. Loffing | (37) |
| NHE3 | Mouse | 1/500 |  | StressMarq | SPC-400 |
| Megalin | Goat |  | 1/2000 | Pr P. Galichon | (38, 39) |
| Uromodulin | Rabbit |  | 1/3000 | Abcam | Ab207170 |
| E-cadherin | Mouse | 1/2000 |  | BD Biosciences | 610404 |
| β-actin | Mouse | 1/10000 | | Sigma | A5441 |
| Aquaporin-2 | mouse |  | 1/3000 | Santa Cruz | sc-515770 |

For quantitative PCR (qPCR), cDNA was combined with 0.5 μM of each primer and SYBR Green Master Mix (cat. no. A25743, Applied Biosystems, Foster City, CA) and reactions were performed in duplicate using the ABI StepOne sequence detection system (Applied Biosystems). Data analysis was conducted with ABI Prism software (Applied Biosystems), utilizing P0 as an internal reference. The fold change in cDNA levels (F) was determined using the equation F = 2^(Ct1−Ct2), where Ct1 and Ct2 represent the cycle thresholds for the experimental and control conditions, respectively.

### Statistics

Results are presented as the mean ± SD from n independent experiments. Prism version 10.4.1 (GraphPad Software, San Diego, CA) was used for statistical analysis. The normality of the data was assessed using the Shapiro-Wilk test. For normally distributed data, statistical differences between two groups were determined using a two-tailed unpaired Student’s t-test and between more than two groups by ANOVA, whereas for non-normally distributed data, the non-parametric Mann-Whitney U-test for two groups or the Kruskal-Wallis test for more than two groups. A p-value < 0.05 was considered significant.

## Results

### Segment-specific expression of claudin-3 along the mouse nephron

To investigate the expression and localization of claudin-3 in the mouse kidney, we first validated the specificity of the claudin-3 antibody. This was achieved by Western blot analysis and immunostaining of kidney tissues obtained from both wild-type and claudin-3 knockout mice (Supplemental Fig. S1), confirming the antibody’s specificity. Immunofluorescence further demonstrated that claudin-3 is localized as expected at the apical pole of renal epithelial cells and displays the typical reticulated pattern of a tight junctional protein.

As an initial step toward elucidating the physiological function of claudin-3 in renal epithelia, we sought to determine its segment-specific distribution along the nephron. Accurate localization of claudin-3 is critical for understanding its role in the structural and functional organization of the kidney. For this purpose, we performed immunofluorescence labeling on paraffin-embedded kidney sections from adult C57BL/6 mice, using a panel of well-characterized segment-specific markers to identify distinct nephron regions. Claudin-3 immunoreactivity was not detected in the glomerulus. In the proximal tubule, identified by apical megalin staining, claudin-3 was likewise absent, indicating that it is not expressed in this segment (Figure 1A). In contrast, claudin-3 expression was clearly detected in the thick ascending limb (TAL) of Henle’s loop, as demonstrated by its colocalization with the TAL-specific marker uromodulin (Figure 1B). Claudin-3 was also expressed in the distal convoluted tubule (DCT), on the basis of its colocalization with the sodium-chloride cotransporter (NCC), a specific marker for this segment (Figure 1C). In the downstream segments of the nephron, including the connecting tubule (CNT) and collecting duct (CD), claudin-3 was found to colocalize with γ-subunit of the epithelial sodium channel (γ-ENaC), which is expressed in both segments (Figure 1D). To specifically label the CD, we used aquaporin-2 (AQP2), a marker specific for the collecting duct principal cells. Colocalization of claudin-3 with AQP2 confirmed its expression along the collecting duct (Figure 1E).

**Figure 1.**
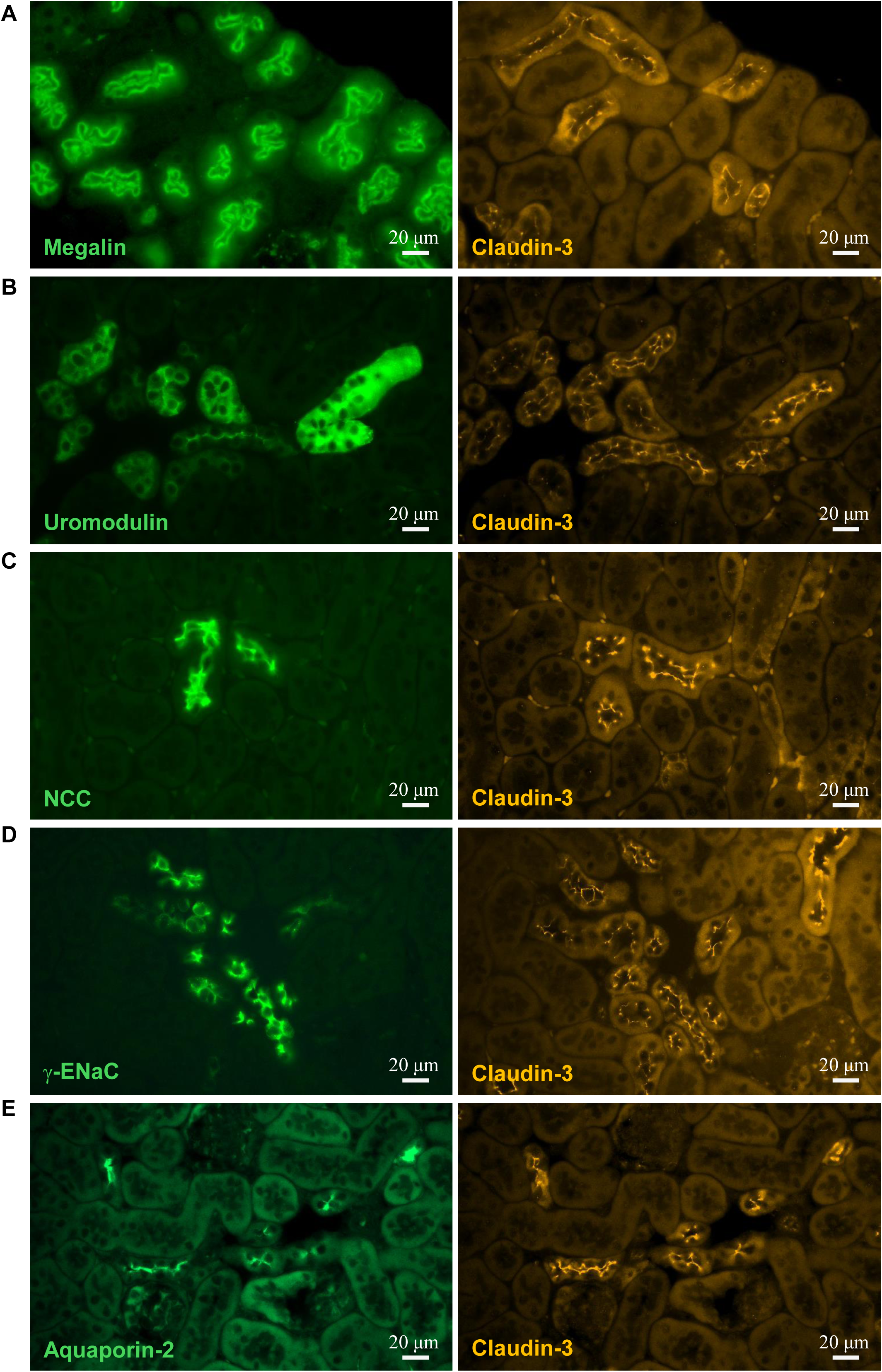
Localization of claudin-3 along the kidney tubule. Representative immunofluorescence staining of serial kidney sections (x40 objective). A section (n) was stained for a specific tubule segment marker (left, green), and the adjacent section (n+1) was stained for claudin-3 (middle, orange). Segment specific markers are megalin for the proximal tubule (A), uromodulin for the thick ascending limb (B), the thiazide-sensitive sodium-chloride cotransporter (NCC) for the distal convoluted tubule (C), the epithelial sodium channel gamma subunit (γ-ENaC) for the connecting tubule and cortical collecting duct tubule (D) and aquaporin-2 (AQP2) for the collecting duct tubule (E).

Taken together, these findings indicate that claudin-3 exhibits a distinct segmental distribution pattern along the nephron. It is selectively expressed in the distal portion of the kidney tubule including TAL, DCT, CNT, and CD, but is absent from both the glomerulus and proximal tubule. This segment-specific expression suggests that claudin-3 may play a functional role in epithelial barrier properties and transepithelial transport in the distal nephron.

### Claudin-3 deletion does not alter basic physiological parameters under basal conditions

To investigate the role of claudin-3 in the kidney, we analyzed a global claudin-3 knockout mouse model (KO) (22). Phenotypic analysis revealed that claudin-3 deficient mice do not exhibit any remarkable phenotype compared to wild-type (WT) controls. They remain fertile and maintain normal body weight (Figure 2A). Similarly, the kidney-to-body weight ratio at 12 weeks of age does not differ significantly between genotypes (Figure 2A). Basic physiological analysis in metabolic cages did not show significant difference in food and water intake or in 24-hour urine output (Figure 2A). Urinary excretion of sodium, potassium, chloride, and basal urine osmolality over 24 hours also remained comparable between KO and WT mice (Figure 2B).

**Figure 2.**
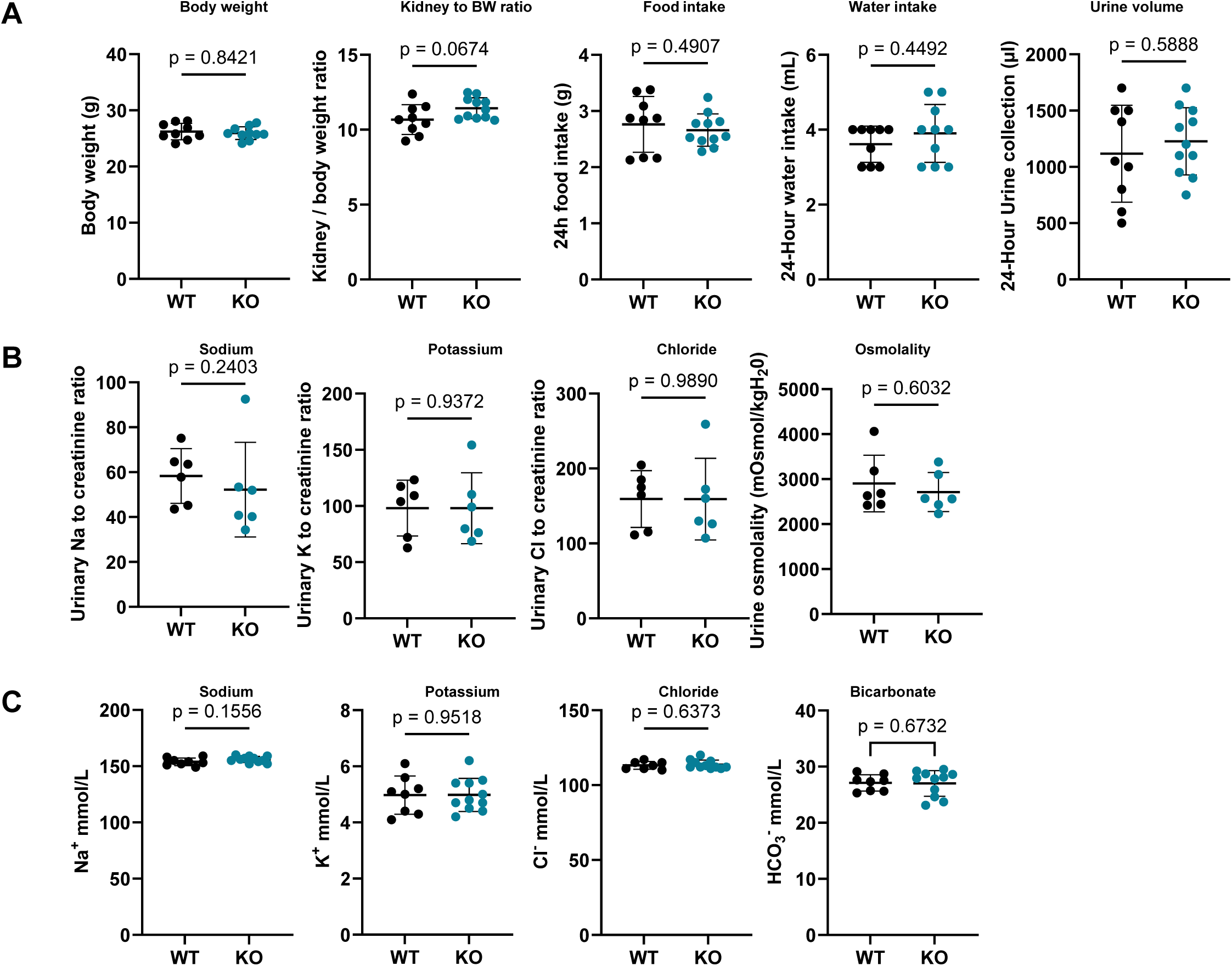
Physiological parameters of claudin-3 deficient mice under a normal sodium diet. Mice were maintained on a normal-sodium diet (0.18%; NSD) and analyzed for various physiological parameters. (A) Mice were first acclimated to the metabolic cages for 24-hour, followed by 24-hour of measurement of physiological parameters, including body weight, kidney to body weight ratio, 24-hour food intake, 24-hour water intake and 24-hour urine volume in WT and claudin-3 KO mice. (B) Urine was collected over a 24-hour period, and urinary Na^+^, Cl^−^, and K^+^ excretion, as well as urine osmolality, were measured in WT and claudin-3-deficient mice. (C) Biological analysis from venous blood showing sodium (Na^+^), potassium (K^+^), chloride (Cl^−^) and bicarbonate (HCO_3_^−^) in WT and claudin-3 KO mice under NSD. Statistical analysis was performed using Mann-Whitney U test. Data are presented as mean ± SD.

Measurements of venous blood gas parameters and plasma electrolyte levels including sodium (Na^+^), potassium (K^+^), chloride (Cl^−^), and bicarbonate (HCO_3_^−^) did not reveal significant differences between the two groups (Figure 2C). These findings indicate that claudin-3 deficiency does not alter overall growth, metabolic parameters, or body fluid homeostasis under baseline conditions.

### Claudin-3 knockout does not alter basal sodium transporters or claudin expression along the nephron

We then investigated the impact of claudin-3 deletion on sodium transport and tight junction proteins along the nephron by measuring the expression of key sodium transporters and claudins in proximal and distal segments using Western blot analysis. We assessed the abundance of NHE3, NKCC2, and NCC (both total and phosphorylated forms), as well as claudin-2 and claudin-10. We found no significant difference between claudin-3 knockout (KO) and wild-type (WT) mice for any of these proteins (Figure 3A-B).

**Figure 3.**
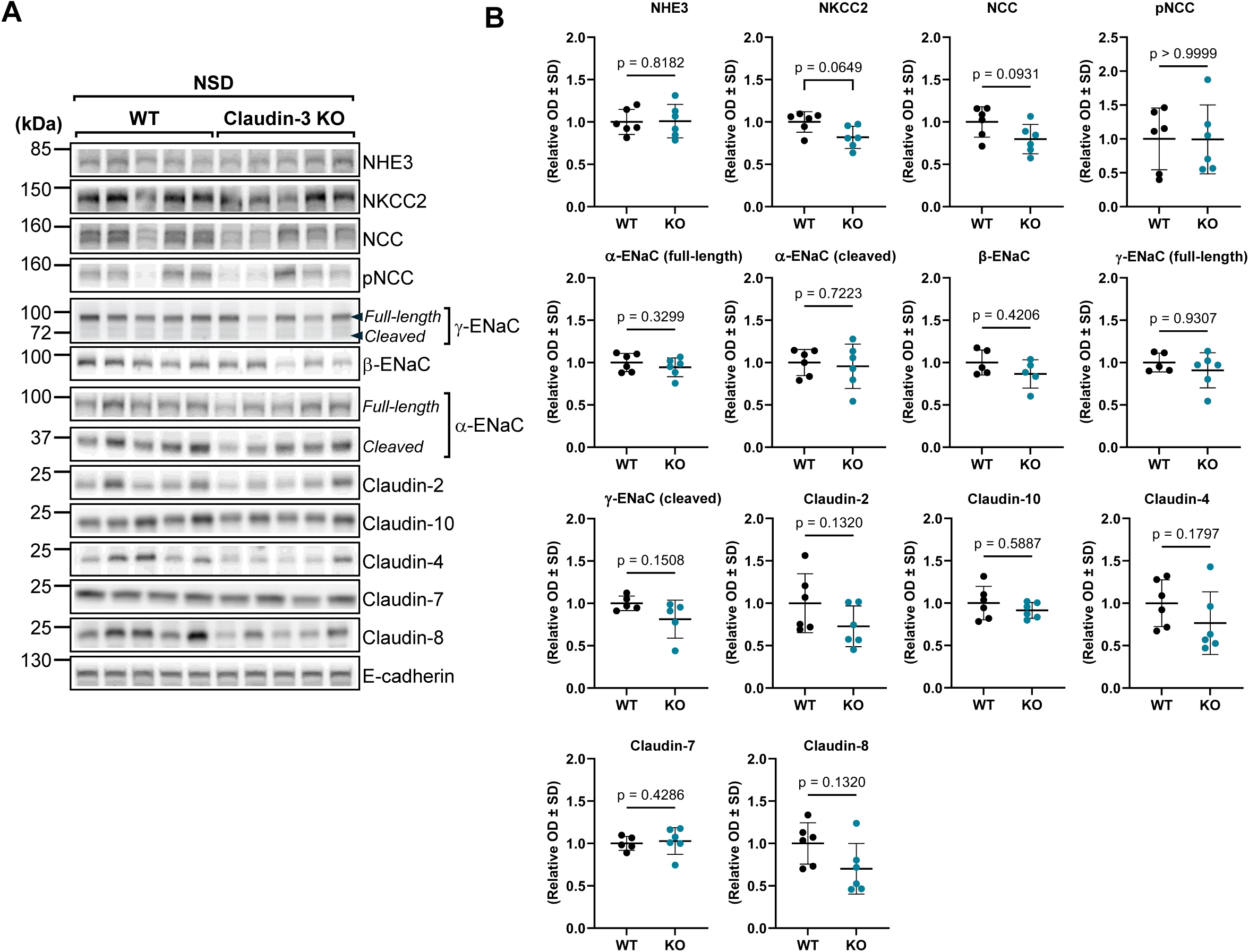
Segmental analysis of sodium transporters and claudins along the nephron in claudin-3 knockout mice under normal sodium intake. Wild-type (WT) and claudin-3 knockout (Claudin-3 KO) mice were maintained on a normal-sodium diet (0.18% Na^+^; NSD). (A) Representative Western blots showing the expression of full-length and cleaved forms of α- and γ-subunits of the epithelial sodium channel (ENaC), β-ENaC, the thiazide-sensitive sodium-chloride cotransporter (NCC; total and phosphorylated), the Na^+^-K^+^-2Cl^−^ cotransporter (NKCC2), and tight junction claudins (claudin-2, claudin-3, claudin-4, claudin-7, claudin-8, and claudin-10) in the renal cortex of wild-type and claudin-3 knockout mice. (B) Densitometric quantification of protein abundance normalized to E-cadherin from five animals per group. Statistical analysis was performed using Mann-Whitney U test. Data are presented as mean ± SD. kDa: kilodaltons; OD: optical density.

The expression of collecting duct-associated claudins (claudin-4, -7, and -8) and epithelial sodium channel (ENaC) subunits (α, β, and γ) also remained unchanged. For α-ENaC and γ-ENaC, both the full-length and active cleaved forms were assessed and no difference between genotypes was observed (Figure 3A-B).

Together, these results demonstrate that claudin-3 deletion does not induce compensatory changes in claudin expression or sodium transport–related proteins along the nephron under basal physiological conditions.

### Aldosterone upregulates claudin-3 in the mouse kidney

Further examining the potential physiological effect of aldosterone on claudin-3 *in vivo*, we analyzed its expression levels in wild type mice subjected to a normal sodium diet (NSD) compared to those fed with a low sodium diet (LSD). Following 7 days of LSD, the expression levels of α-ENaC were assessed in the renal cortex by Western blot analysis. Both the full-length and cleaved forms of α-ENaC were examined. While the cleaved form was significantly upregulated, the full-length form showed a trend toward increased expression (Figure 4A-B). These results confirm that LSD leads to an increase in endogenous aldosterone biological activity. Interestingly, claudin-3 protein abundance, assessed by both Western blot and immunofluorescence, was more abundant in the kidney of LSD-fed mice (Figure 4A-C). Notably, our previous study demonstrated that LSD induces claudin-8 expression in mice while does not affect other collecting duct claudins, including claudin-4 and claudin-7 (25).

**Figure 4.**
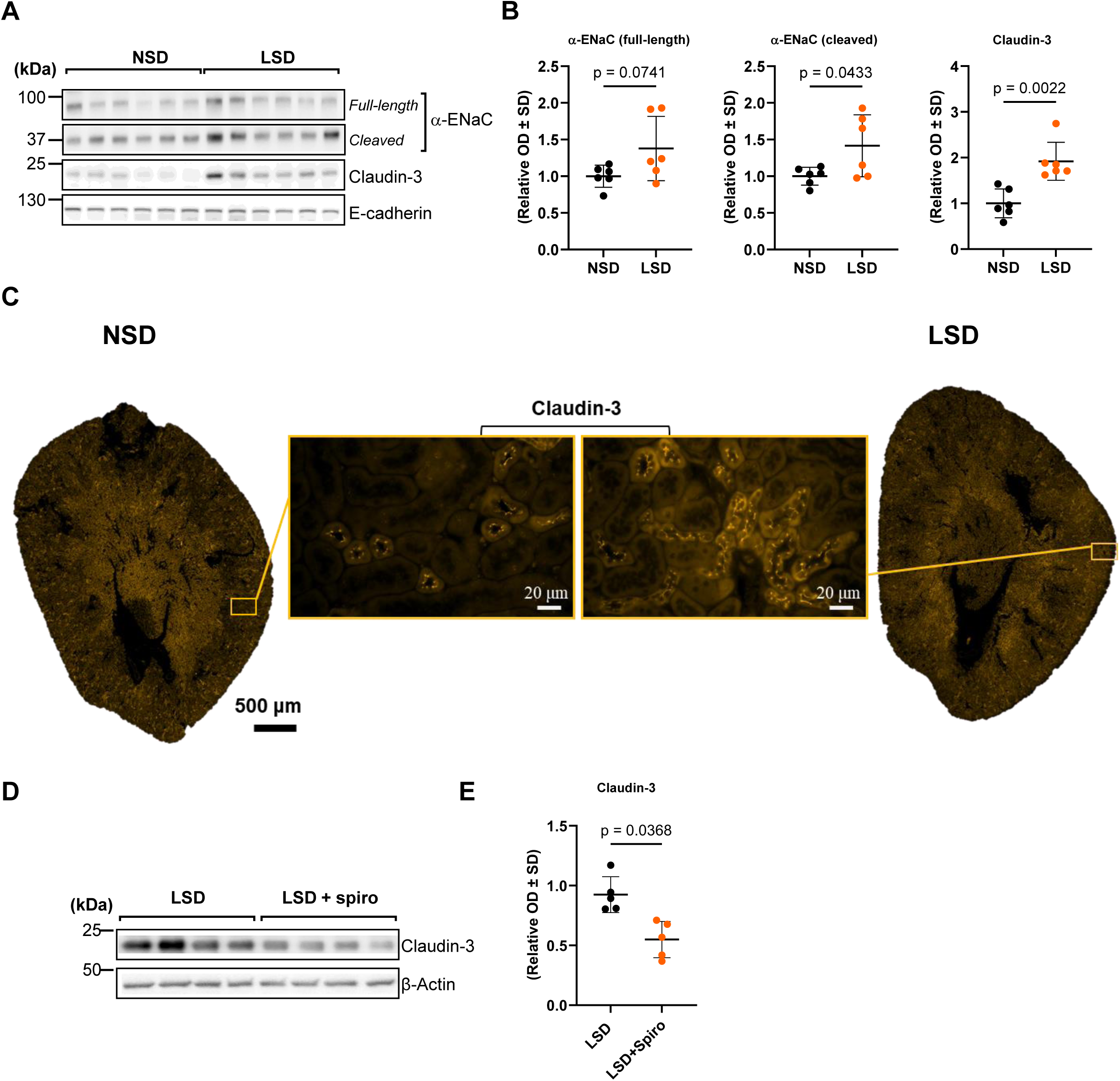
Aldosterone regulates claudin-3 expression in the mouse kidney. C57BL/6 mice were maintained on either a normal-sodium (Na^+^) diet (0.18%; NSD) or a low-sodium diet (0.01%; LSD) for seven days. (A) Western blot analysis illustrating the impact of dietary sodium intake on the abundance of claudin-3 and α-epithelial sodium channel (α-ENaC) in the kidney cortex. Representative blots from six animals per experimental group are shown. (B) Relative densitometric quantification of immunoblots from the kidney cortex, with E-cadherin used as a loading control. (C) Representative immunofluorescence imaging of claudin-3 in kidney transverse sections from NSD and LSD mice (x20 objective), shown in orange. Insets of each image are shown with a x40 objective. (D) Effect of mineralocorticoid receptor blockade on claudin-3 expression. Mice were fed an LSD (0.01%) and treated or not with spironolactone (0.35 mg/100 g body weight/day) for one week before kidney cortex protein extraction. (D) Western blot analysis showing the effect of spironolactone on claudin-3 expression in the kidney cortex of four animals per experimental group. (E) Densitometric quantification of immunoblots from the kidney cortex of six animals, with β-actin used as a loading control. Statistical analysis was performed using Mann-Whitney U test. Data are presented as mean ± SD. kDa: kilodaltons; OD: optical density.

To further confirm the role of aldosterone in the regulation of claudin-3 expression, LSD-fed mice were treated for 7 days with spironolactone, a mineralocorticoid receptor antagonist. The blockade of MR with spironolactone effectively inhibited the LSD-induced increase in claudin-3 protein abundance, demonstrating that this effect is dependent on activation of the MR by endogenous aldosterone (Fig. 4D-E).

To validate these findings *in vitro,* we used mCCD_cl1_ cells, a model of cortical collecting duct principal cells (26). Treatment with aldosterone (10^−^⁶ M, 24 h) significantly increased claudin-3 expression at both protein and mRNA levels, as shown by Western blot, immunofluorescence, and qPCR analyses (Supplemental Fig. S2A-D). Supporting this conclusion, *in silico* analysis using the Eukaryotic Promoter Database (https://epd.epfl.ch) identified multiple MR-binding sites in the promoter region of the mouse claudin-3 gene (Supplemental Fig. S2E), consistent with genomic effect of aldosterone mediated by the MR. Notably, our previous findings indicated that aldosterone increases claudin-8 expression in the collecting duct but does not alter the expression of claudin-4 and claudin-7, two other claudins expressed in this segment (25).

Functionally, overexpression of claudin-3 in mCCD_cl1_ cells reduced paracellular permeability to sodium and chloride (Supplemental Fig. S3A–D), while silencing claudin-3 increased permeability to both ions (Supplemental Fig. S4A–D). These results further demonstrate that claudin-3 is induced by aldosterone and indicates that it functions as a paracellular sodium-chloride barrier in CD principal cells.

Together, these findings show that fluctuations in endogenous aldosterone levels regulate claudin-3 abundance in the mouse kidney and identify claudin-3 as a functional target of aldosterone in the collecting duct epithelium.

### Low salt diet induces segment-specific compensatory upregulation of ENaC subunits and claudins in claudin-3-deficient mice

Since we observed an induction of claudin-3 expression by aldosterone, we investigated the impact of claudin-3 deletion on sodium transport under sodium-restricted conditions to reveal its functional effect *in vivo*. Claudin-3 knockout (KO) and wild-type (WT) mice were fed a low-salt diet (LSD) for 7 days. LSD induced a significant increase in both cleaved and full-length forms of α-ENaC and γ-ENaC protein levels in the renal cortex of KO compared to WT mice, as shown by Western blot analysis (Figure 5A-B). The upregulation of γ-ENaC was also confirmed by immunofluorescence (Figure 5C). In contrast, β-ENaC abundance remained unchanged (Figure 5A-B).

**Figure 5.**
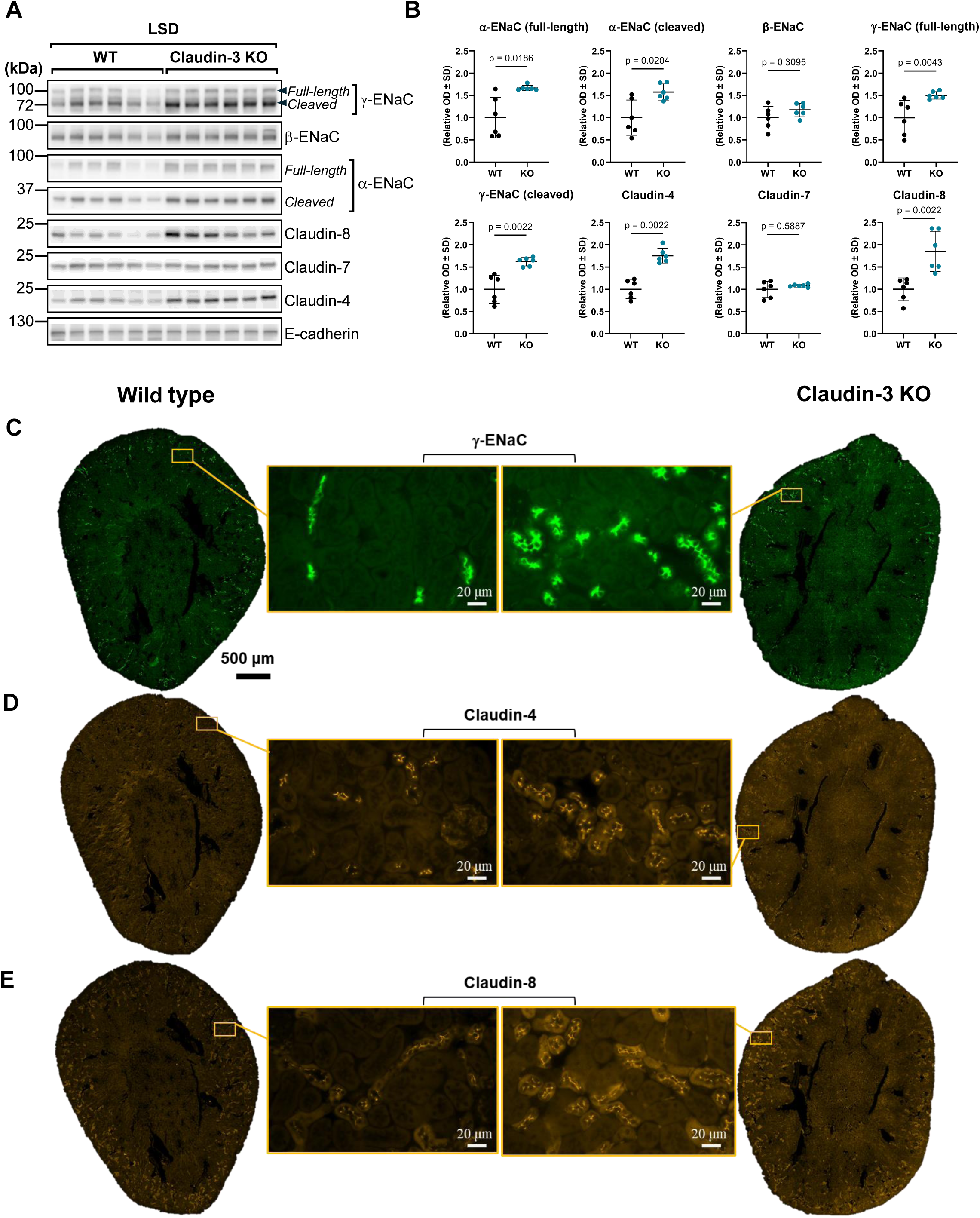
Compensatory upregulation of ENaC subunits and collecting duct claudins in claudin-3 deficient mice under a low sodium diet. Wild-type (WT) and claudin-3 knockout (Claudin-3 KO) mice were maintained on a low-sodium diet (0.01%; LSD) for seven days. (A) Western blot analysis showing the effect of dietary sodium intake on the abundance of α-epithelial sodium channel (α-ENaC), γ-epithelial sodium channel (γ-ENaC), claudin-4, claudin-7, and claudin-8 in the kidney cortex. (B) Relative densitometric quantification of immunoblots from the kidney cortex of six animals, with E-cadherin used as a loading control. (C-E) Representative immunofluorescence images of γ-ENaC (C), claudin-4 (D), and claudin-8 (E) in control and claudin-3 KO mice (x20 objective). Insets of each image are shown with a x40 objective. Statistical analysis was performed using Mann-Whitney U test. Data are presented as mean ± SD. kDa: kilodaltons; OD: optical density.

We also assessed the expression of claudins expressed in the collecting duct, given their role in tight junction integrity and potential involvement in compensatory epithelial responses. Claudin-4 and claudin-8 protein abundance was increased under LSD in claudin-3 KO mice, as shown by Western blot (Figure 5A-B) and immunofluorescence (Figure 5A-B). In contrast, claudin-7 levels remained unchanged (Figure 5D-E), indicating a selective regulation of claudin expression in response to sodium restriction combined with claudin-3 deficiency.

Beyond the collecting duct, we examined other nephron segments where claudin-3 is also expressed. We analyzed total and phosphorylated NCC, NKCC2, and claudin-10 protein abundance by Western blot (Figure 6A-B). No significant difference in total or phosphorylated NCC levels between WT and KO mice was observed. NKCC2 exhibited a trend toward increased expression in KO mice, but the difference did not reach statistical significance. Importantly, Western-blot analysis showed that claudin-10 expression was significantly increased in KO mice. Immunofluorescence analysis revealed that this upregulation of claudin-10 took place along the thick ascending limb of Henle’s loop (Figure 6A-C).

**Figure 6.**
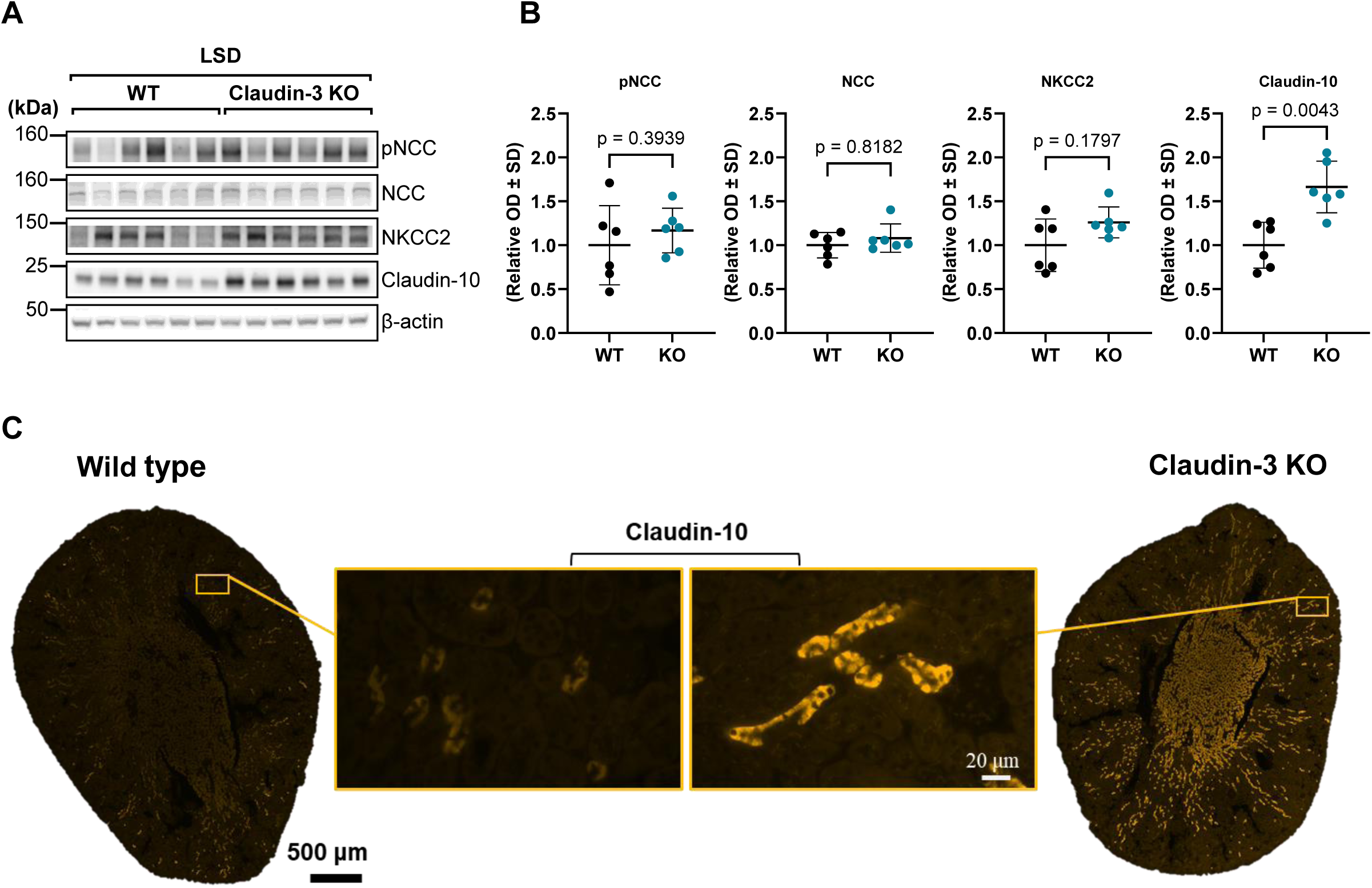
Differential regulation of sodium transporters and claudin-10 in claudin-3 deficient mice under a low sodium diet. Wild-type (WT) and claudin-3 knockout (Claudin-3 KO) mice were maintained under a low-sodium diet (0.01%; LSD) for seven days. (A) Western blot analysis showing the effect of sodium restriction on total NCC, phosphorylated NCC (p-NCC), NKCC2, and claudin-10 in kidney cortex lysates. (B) Densitometric quantification of protein abundance of immunoblots from the kidney cortex of six animals, normalized to β-actin. (C) Representative immunofluorescence images of claudin-10 in WT and claudin-3 KO mice (shown in orange, ×20 objective). Insets show higher magnification (×40 objective). Statistical comparisons were performed using two-tailed t-tests. kDa, kilodaltons; OD, optical density.

Together, these results demonstrate that in the absence of claudin-3, a low-salt diet triggers segment-specific compensatory mechanisms, including the upregulation of ENaC subunits in association with a specific set of claudins along the collecting duct and upstream distal nephron segments, likely contributing to the maintenance of sodium balance and epithelial barrier function.

### Mechanisms of adaptation to low-sodium challenge in claudin-3-deficient mice

To investigate whether aldosterone-mineralocorticoid receptor (MR) signaling mediates the compensatory response to a low-sodium diet (LSD) in claudin-3 knockout (KO) mice, we treated WT and KO animals with spironolactone, a selective MR antagonist, under LSD conditions. Western blot analysis shows that MR blockade did not abolish the upregulation of both cleaved and full-length α-ENaC in KO mice relative to WT (Figure 7A-B). In contrast, the increase in full-length γ-ENaC no longer reaches statistical significance under MR antagonism, although cleaved γ-ENaC remains elevated (Figure 7A-B). In addition, claudin-4, claudin-8 and claudin-10 remained significantly induced in KO mice under MR antagonism (Figure 7A-B). These result suggest that aldosterone does not mediate the adaptations of transcellular and paracellular sodium transport that take place under low sodium diet in claudin-3 deficient mice.

**Figure 7.**
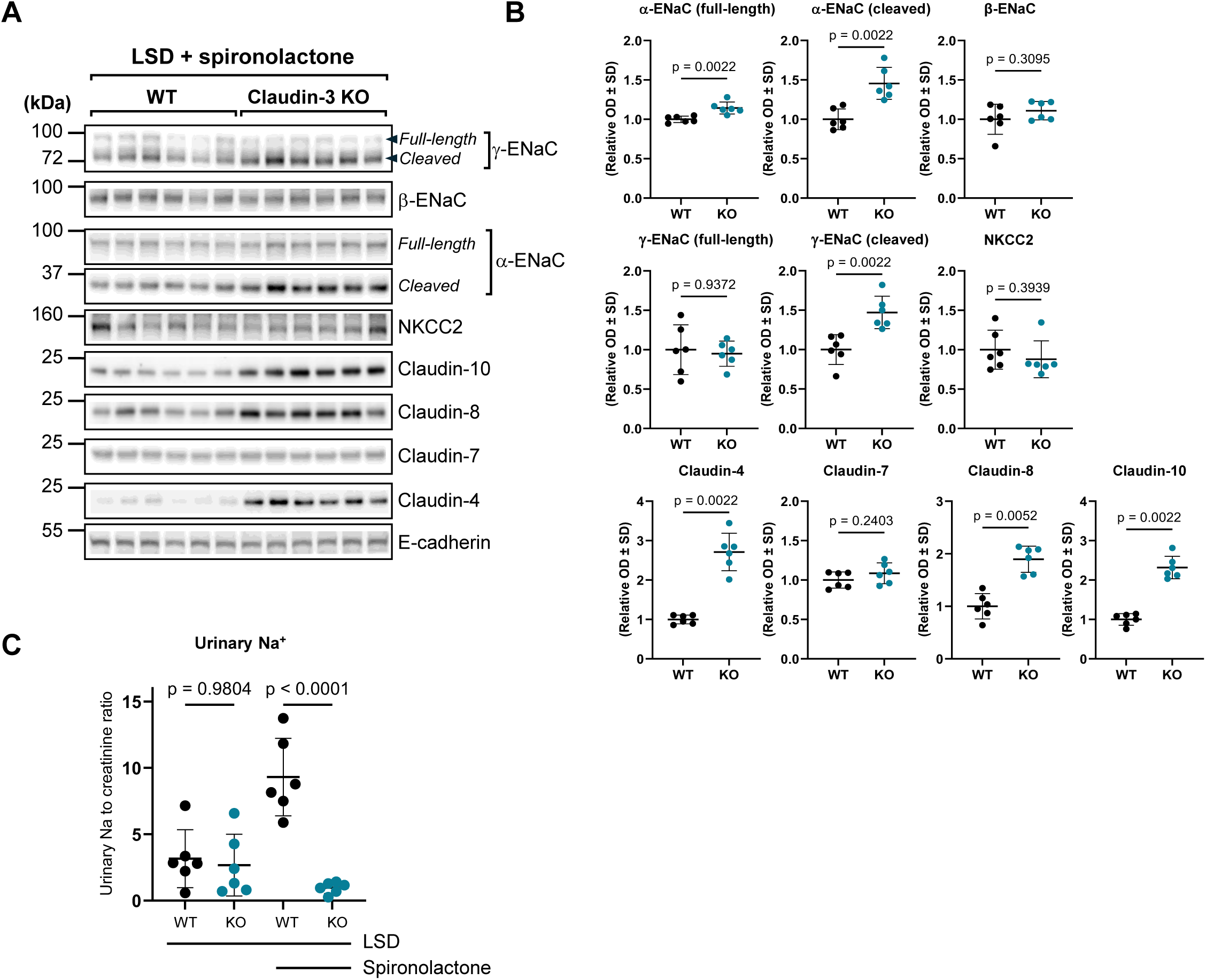
Effect of spironolactone in WT claudin-3 deficient mice. Wild-type (WT) and claudin-3 knockout (Claudin-3 KO) mice were fed a low-sodium diet (0.01%, LSD) and treated with spironolactone (0.35 mg/100 g body weight/day) for 7 days before kidney cortex protein extraction. (A) Representative Western blots showing the abundance of ENaC subunits (α, β and γ), NKCC2, claudin-4, claudin-7, claudin-8 and claudin-10 protein abundance in the kidney cortex of six animals per experimental group. (B) Densitometric quantification of immunoblots from (A), normalized to E-cadherin as a loading control. Data are presented as mean ± SD. (C) Urinary Na^+^ excretion over a 24-hour period measured in WT and claudin-3 KO mice. Statistical analysis was performed using Mann-Whitney U test (B) and ordinary one-way ANOVA (C), a p value < 0.05 was considered significant. kDa, kilodaltons ; OD, optical density.

Urinary sodium measurements (Figure 7C) show that WT mice treated with spironolactone excreted significantly more sodium compared to untreated controls, confirming the efficacy of MR blockade in inhibiting sodium reabsorption. In contrast, KO mice treated with spironolactone did not exhibit increased sodium excretion, supporting the Western blot findings that compensatory sodium transport regulation in KO mice persists despite MR antagonism.

These findings demonstrate that the compensatory upregulation of ENaC subunits and multiple claudins in claudin-3-deficient mice on a low-sodium diet is not exclusively mediated by MR activation. The persistence of α-ENaC, cleaved γ-ENaC, claudin-4, claudin-8 and claudin-10 induction under MR blockade strongly suggests an aldosterone-independent compensatory mechanism aimed at preserving sodium balance.

## Discussion

Claudin-3, a tight junction protein, is critically involved in renal function through its role in paracellular permeability. Here, we aimed to characterize its *in vivo* distribution along the nephron, determine its functional significance in sodium handling, and assess its potential regulation by aldosterone.

Our segment-specific mapping of claudin-3 along the mouse nephron consolidates and extends previous observations on claudin distribution. Distinct claudin isoforms confer unique paracellular properties to different nephron segments: claudin-3 is enriched in the distal convoluted tubule, connecting tubule, and collecting duct alongside claudins-4, -7, and -8 (27). It is also expressed in the thick ascending limb of Henle’s loop where it contributes to barrier tightening (28, 29). It should be mentioned that in this later segment, claudin-3 expression is restricted to junctions containing claudin-16 and claudin-19 (30). By definitively placing claudin-3 at tight junctions in the TAL, DCT, CNT and CD, and demonstrating its absence from proximal regions, our findings provide the anatomical basis for future functional studies on how claudin-3 contributes to segment-specific barrier properties and electrolyte handling in the distal nephron.

Aldosterone is well established as the principal hormonal driver of Na^+^ reabsorption and K^+^ secretion in the aldosterone-sensitive distal nephron (ASDN), which encompasses the late distal convoluted tubule, connecting tubule and cortical collecting duct (31, 32). In this context, our findings position claudin-3 as a novel aldosterone target whose upregulation reinforces paracellular barrier function in the collecting duct. *In vitro*, aldosterone treatment of mCCD_cl1_ cells increases claudin-3 abundance and enhances its lateral membrane localization. Claudin-3 upregulation is associated with a decrease in Na^+^ and Cl^−^ permeability, consistent with observations in MDCK cells reported by Fromm’s group (15). Extending these findings to an *in vivo* context, we demonstrate that a low sodium diet which is established to increase endogenous aldosterone levels (33, 34) induces a significant increase in claudin-3 abundance within the renal cortex of mice. This induction is abolished by administration of spironolactone, a mineralocorticoid receptor antagonist, indicating that claudin-3 upregulation in response to low-sodium diet is MR-dependent. Importantly, in a previous study (25), we showed that aldosterone similarly upregulates claudin-8 in collecting duct epithelia. Collectively, these results support a model wherein aldosterone, acting through MR-mediated transcriptional mechanisms, enhances tight junction barrier function *via* selective induction of claudin-3 and claudin-8, thereby minimizing paracellular back-flux of reabsorbed ions. This paracellular reinforcement likely complements canonical stimulation of ENaC to maximize net sodium conservation under conditions of salt depletion and underscores the importance of tight junction remodeling as an underappreciated yet critical mechanism of mineralocorticoid-driven electrolyte homeostasis.

Our experiments in claudin-3 knockout mice demonstrate that under baseline conditions, these animals display no detectable alterations in renal function, electrolyte balance, changes in sodium transport-related proteins, including other claudins and ENaC subunits, along the nephron. This observation suggests that claudin-3 is dispensable under physiological conditions, where the collecting duct is presumably minimally active due to relatively high dietary sodium intake. This observation aligns with predictions from mathematical models (33, 35) which indicate that sodium reabsorption in the collecting duct contributes minimally to overall sodium handling during normal sodium intake. Furthermore, these models predict that the functional importance of collecting duct increases substantially under sodium-restricted conditions. Consistent with this, we show that claudin-3 knockout mice exhibit a pronounced compensatory response when subjected to a low-sodium diet. This response includes significant upregulation of α- and γ-ENaC subunits and increased expression of claudin-4, claudin-8, and claudin-10. One could speculate that increased ENaC and claudin-4 and -8 abundance would compensate for the paracellular sodium leakage generated by the absence of claudin-3 along the ASDN. In addition, increased claudin 10 expression in the TAL would increase paracellular sodium reabsorption along the lumen positive electrical gradient. Importantly, these adaptations are not prevented by mineralocorticoid receptor blockade with spironolactone, suggesting that alternative mechanisms contribute to maintaining sodium homeostasis in the absence of claudin-3.

Our results expand the understanding of the role of aldosterone in renal sodium handling. Beyond its well-established effects on ENaC- and Na,K-ATPase-mediated transcellular sodium transport (11–13, 36), aldosterone modulates the paracellular pathway through selective upregulation of claudin-3 in addition to claudin-8 (25). This dual mechanism enhancing sodium reabsorption via transcellular route and preventing the back-flux of ions to the lumen via the paracellular route appears critical to increase the efficiency of the reabsorption process. This mechanism might be crucial to maintain electrolyte balance, especially under conditions of low dietary salt intake. Future studies should be designed to elucidate the precise processes governing claudin-3 regulation and explore whether similar mechanisms operate in human kidney epithelia, which may have significant implications for treating disorders such as hypertension and salt-sensitive renal disease.

Collectively, these findings support the concept that modulation of paracellular permeability via tight junction proteins represents an underappreciated but physiologically significant mechanism in distal nephron adaptation to sodium depletion. A model summarizing these findings in both WT and claudin-3 knockout mice is shown in Figure 8.

**Figure 8.**
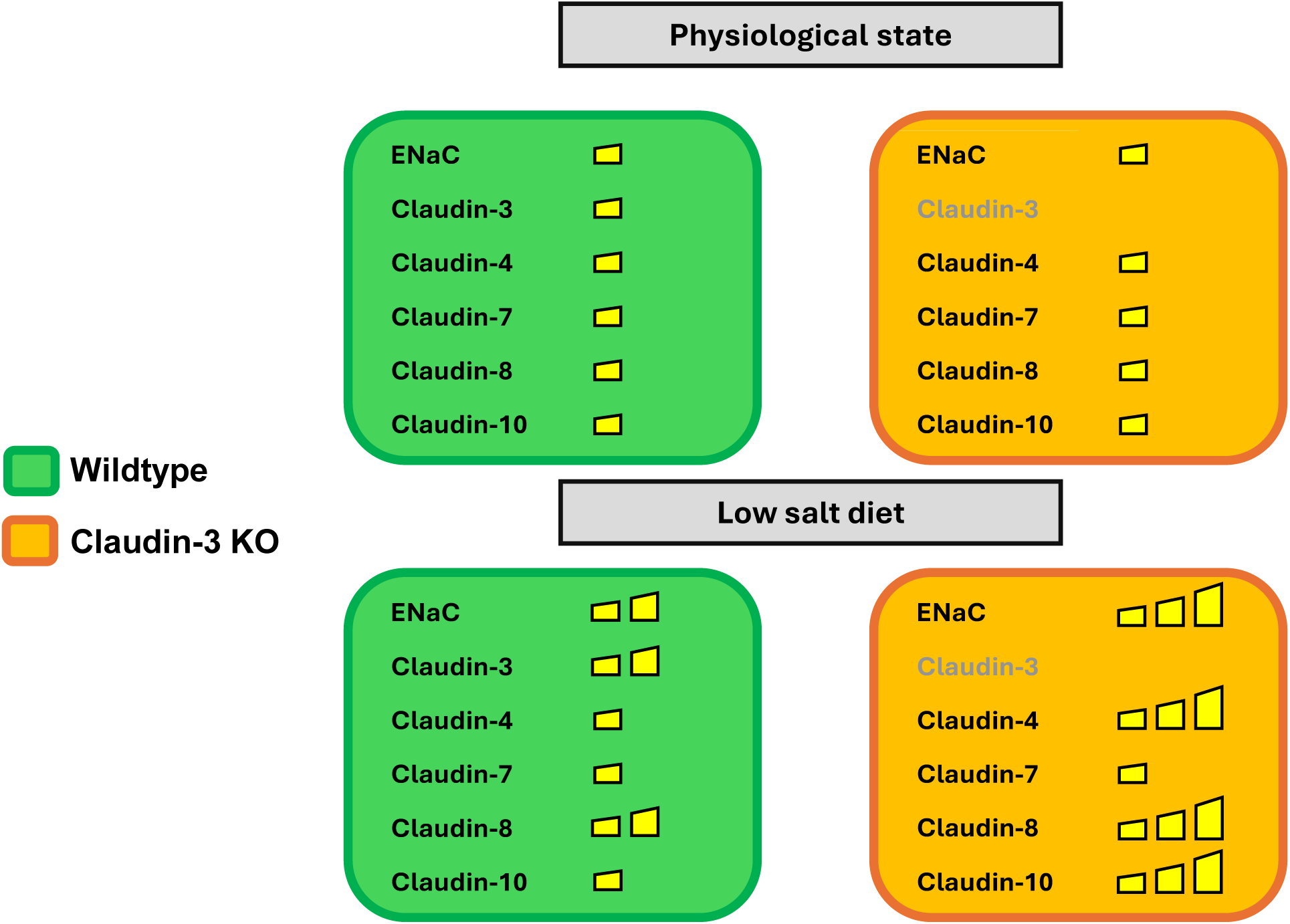
Proposed model of low-sodium-induced adaptations in ENaC and tight junction proteins in claudin-3 knockout and wild-type mice. Under physiological conditions, ENaC expression and the levels of claudin-4, claudin-7, claudin-8, and claudin-10 in the distal nephron are comparable between wild-type (green boxes) and claudin-3 knockout (orange boxes) mice. Under a low-sodium diet, ENaC expression increases in both groups, with a significantly greater upregulation in knockout mice. Wild-type mice respond by upregulating claudin-3 and claudin-8, whereas knockout mice adapt to the absence of claudin-3 by increasing the abundance of claudin-4, claudin-8, and claudin-10 as a compensatory mechanism. Despite these changes, claudin-7 levels remain stable under both physiological and low-sodium diet conditions in WT and KO mice. These adaptations help maintain epithelial barrier integrity and sodium homeostasis despite the absence of claudin-3.

## Perspectives and Significance

Our findings show that aldosterone regulates Na^+^ transport in the collecting duct by enhancing ENaC-mediated reabsorption and modulating tight junctions composition through claudin-3 upregulation, reducing paracellular Na^+^ back-leak. The compensatory adaptations found in claudin-3-deficient mice highlight the interplay between transcellular and paracellular transport. This study underscores the dynamic regulation of the junctional complex and the need for further investigation into hormonal control of the paracellular pathway.

## Authors contributions

E.F and A.S. designed the study, A.S., A.C., S.J., and A.P. carried out experiments, N.L., developed the automatic quantification method for immunofluorescence images, F.B., D.S. and M.F. provided materials and technical assistance, A.S. and A.C. analyzed the data, A.S. and A.C. made the figures, A.S. and E.F. drafted and revised the paper, all authors approved the final version of the manuscript.

## Conflict of interest

The authors declare no conflicts of interest.

## Acknowledgements

This work was supported by the National Center of Competence in Research Kidney Control of Homeostasis and a Swiss National Science Foundation grant 31003A_156736/1 and 207441 to EF and AS. We thank Pr Johannes Loffing (Institute of Anatomy, University of Zürich) and Pr Galichon (INSERM, Paris, France) for kindly providing α-ENaC subunit and megalin antibodies, respectively.

---

**Supplementary Figure 1.**
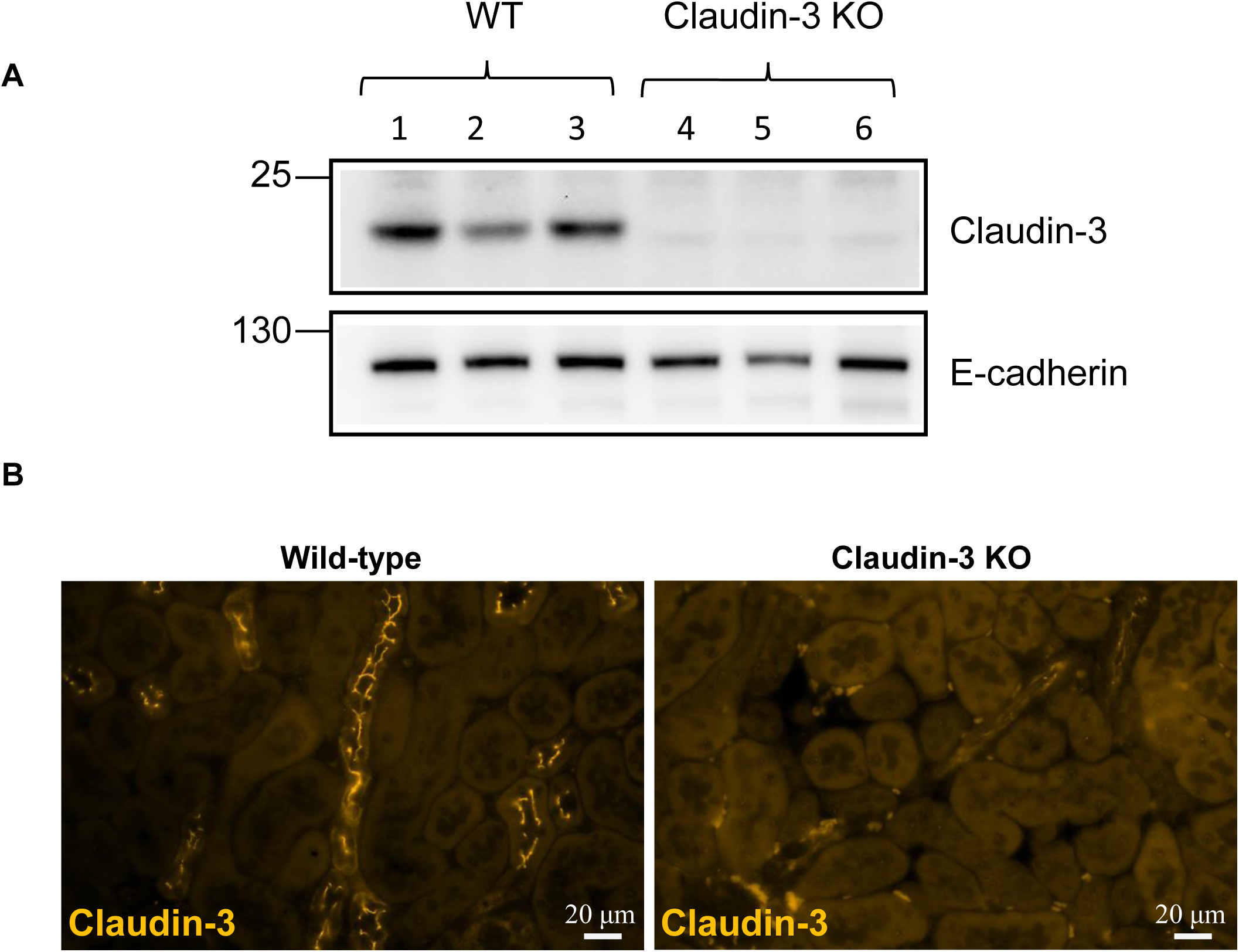
Validation of claudin-3 antibody specificity. The specificity of the claudin-3 antibody was validated using both Western blot (A) and immunofluorescence (B) analyses in wild-type (WT) and claudin-3 knockout (Claudin-3 KO) mouse kidney tissues. (A) Western blot analysis demonstrates a specific band corresponding to claudin-3 at the expected molecular weight in WT tissue, which is absent in claudin-3 KO samples, confirming antibody specificity. (B) Representative immunofluorescence images show claudin-3 localization in WT kidneys (signal in orange; ×40 objective), with no detectable signal in claudin-3 KO tissue, further confirming specificity.

**Supplementary Figure 2.**
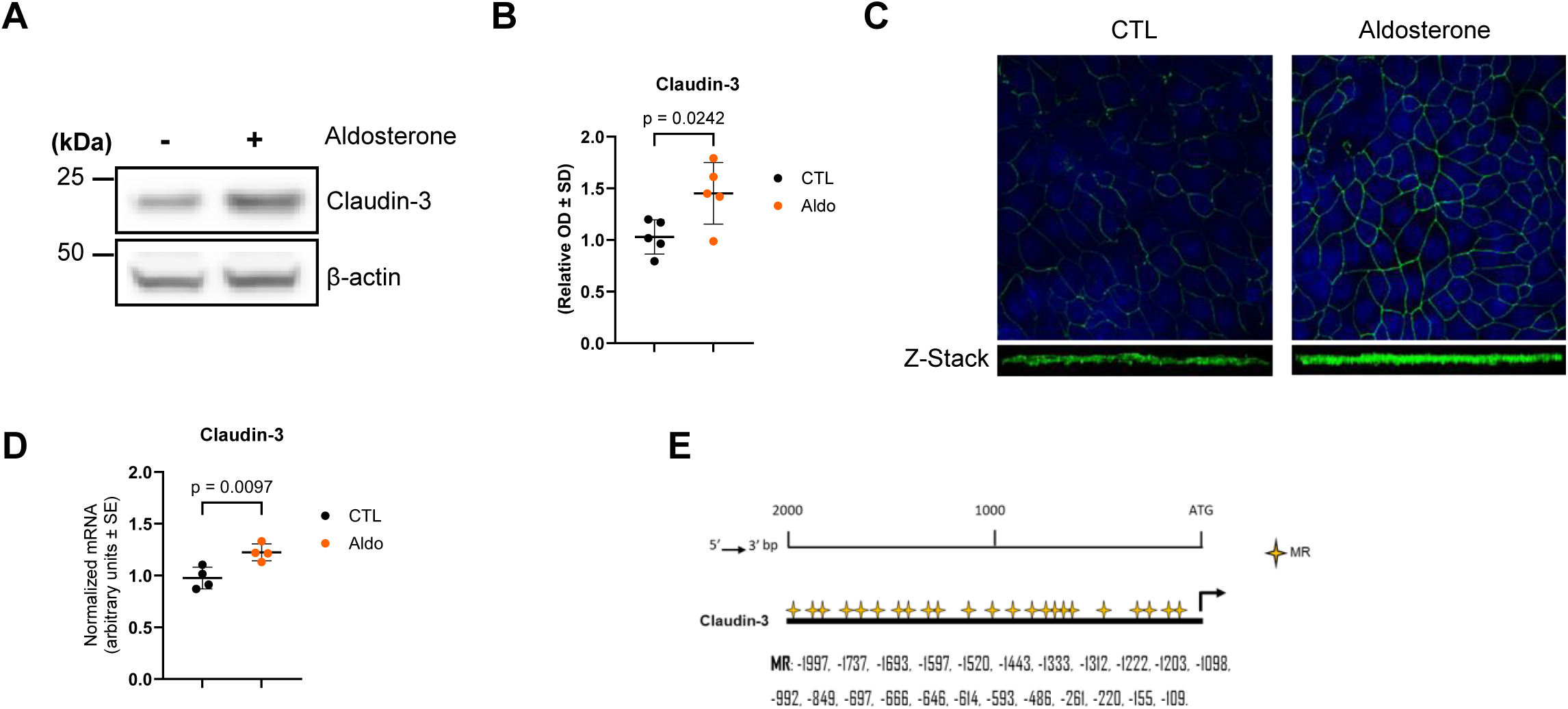
Aldosterone modulates the expression of claudin-3 in cultured mouse collecting duct principal cells. mCCD_cl1_ cells were grown to confluence on filters and treated or not treated with 10^−^⁶ M aldosterone (Aldo) for 24 h. (A) Representative immunoblots showing the effect of aldosterone (aldo) treatment on claudin-3 protein abundance. β-Actin was used as a loading control. (B) Densitometric quantification of immunoblots from (A), from at least 4 independent experiments. (C) Immunofluorescence staining of claudin-3 in confluent monolayers treated or not with aldosterone. DAPI is used for nuclear counterstaining. Lower panels show optical sections from Z-stacks. (D) Claudin-3 mRNA levels assessed by real-time PCR. Results are means ± SD from 4 independent experiments. (E) Diagram of the 2-kb putative promoter region of the mouse *Claudin-3* gene, showing predicted binding sites for the mineralocorticoid receptor (MR). Predictions were obtained using the Eukaryotic Promoter Database (https://epd.epfl.ch). Statistical analysis was performed by a Mann-Whitney U test for comparisons between two groups; *p<0.05,**p<0.01,***p<0.001. Ctl, control; kDa, kilodaltons ; OD, optical density.

**Supplementary Figure 3.**
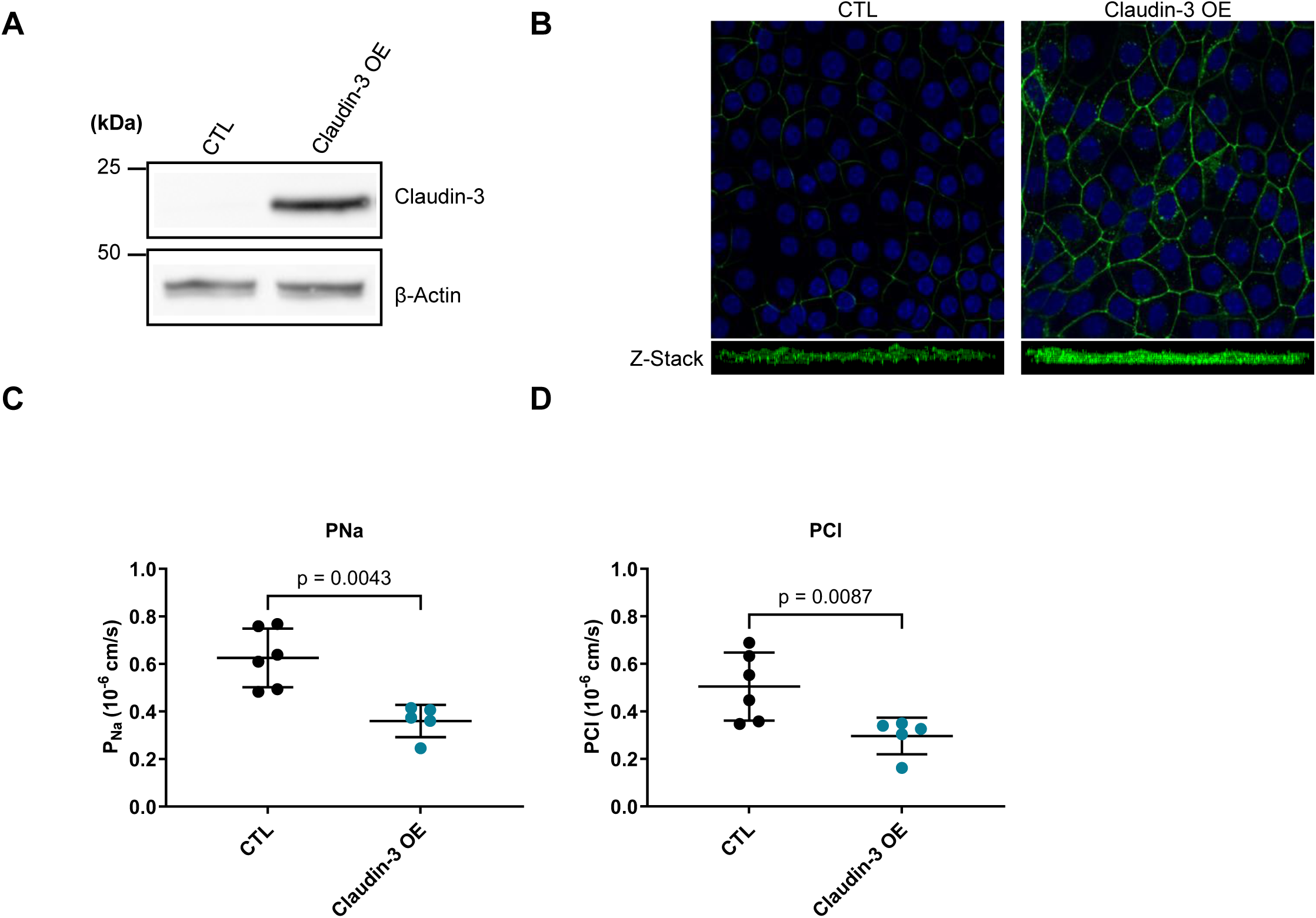
Claudin-3 overexpression decreases paracellular permeability to both Na^+^ and Cl^−^. mCCD_cl1_ cells were transduced with lentiviruses carrying either an empty vector, wild-type green fluorescent protein, or a construct encoding wild-type mouse claudin-3 (Claudin-3 OE) and grown to confluence on filters. (A) Representative immunoblots showing the validation of claudin-3 overexpression. β-actin was used as a loading control. (B) Immunofluorescence staining of claudin-3 in confluent monolayers. DAPI is used for nuclear counterstaining. Lower panels are optical sections from Z-stacks. mCCD_cl1_ cells transduced with empty lentiviruses served as controls in panels A and B. (C-D) Absolute permeabilities for Cl^−^ and Na^+^ calculated from relative ion permeability ratios and transepithelial resistance. Cells transduced with wild-type GFP served as controls in panels C and D. Results are means ± SD from at least four independent experiments. Results are means ± SD from 4 independent experiments. Statistical analysis was performed by a Mann-Whitney U test for comparisons between two groups; *p<0.05,**p<0.01,***p<0.001. Ctl, control; kDa, kilodaltons.

**Supplementary Figure 4.**
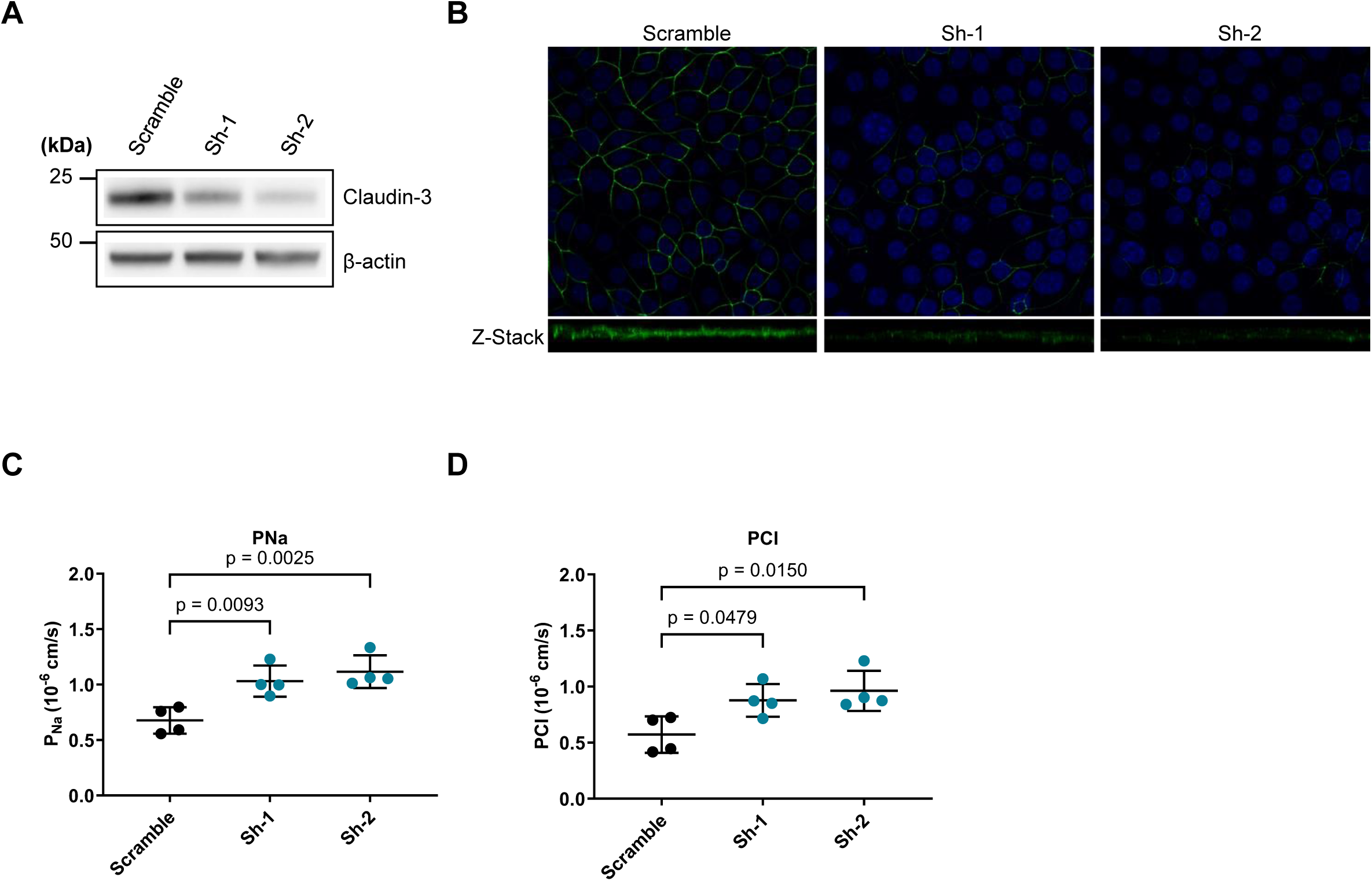
Claudin-3 silencing increases paracellular permeability to both Na^+^ and Cl^−^. mCCD_cl1_ cells transduced with lentiviruses encoding either scramble shRNA (scramble) or shRNA targeting mouse claudin-3 (Sh-1 and Sh-2) were grown to confluence on filters. (A) Representative immunoblots showing the validation of the silencing of claudin-3. β-Actin was used as a loading control. (B) Immunofluorescence staining of claudin-3 in confluent monolayers. DAPI is used for nuclear counterstaining. Lower part of each panel is an optical section obtained from Z-stack. (C-D) Absolute permeabilities of Na^+^ and Cl^-^ calculated from relative ion permeability ratios and the transepithelial resistance measured during the same experiment. Results are means ± SD from at least four independent experiments. Statistical analysis was performed by a Mann-Whitney U test and one-way ANOVA for comparisons between more than two groups; *p<0.05,**p<0.01,***p<0.001. kDa, kilodaltons.

## Notes

### Competing Interest Statement

The authors have declared no competing interest.

